# Structural basis of TFIIIC-dependent RNA Polymerase III transcription initiation

**DOI:** 10.1101/2023.05.16.540967

**Authors:** Anna Talyzina, Yan Han, Chiranjib Banerjee, Susan Fishbain, Alexis Reyes, Reza Vafabakhsh, Yuan He

**Affiliations:** Department of Molecular Biosciences, Northwestern University, Evanston, IL, United States; Interdisciplinary Biological Sciences Program, Northwestern University, Evanston, IL, United States; Chemistry of Life Processes Institute, Northwestern University, Evanston, IL, United States; Robert H. Lurie Comprehensive Cancer Center of Northwestern University, Northwestern University, Chicago, IL, United States; Lead contact

**Keywords:** Transcription, RNA Polymerase III, TFIIIC, TFIIIA, TFIIIB, cryo-EM, FRET

## Abstract

RNA Polymerase III (Pol III) is responsible for transcribing 5S ribosomal RNA (5S rRNA), tRNAs, and other short non-coding RNAs. Its recruitment to the 5S rRNA promoter requires transcription factors TFIIIA, TFIIIC, and TFIIIB. Here we use cryo-electron microscopy to visualize the *S. cerevisiae* complex of TFIIIA and TFIIIC bound to the promoter. Brf1-TBP binding further stabilizes the DNA, resulting in the full-length 5S rRNA gene wrapping around the complex. Our smFRET study reveals that the DNA undergoes both sharp bending and partial dissociation on a slow timescale, consistent with the model predicted from our cryo-EM results. Our findings provide new insights into the mechanism of how the transcription initiation complex assembles on the 5S rRNA promoter, a crucial step in Pol III transcription regulation.

## INTRODUCTION

RNA Polymerase III (Pol III) transcribes various types of short, non-coding, and abundant RNAs from three types of promoters. The Type I promoter is found in the 5S ribosomal RNA (rRNA) gene, the Type II promoter is associated with transfer RNA (tRNA) genes, and the Type III promoter is used in U6 small nuclear RNA (snRNA) genes and others ^1-3^. Both Type I and Type II promoters contain internal control regions (ICRs) in the gene body ^4,5^. The ICRs of the Type I promoter harbor an A-box, an intermediate element (IE), and a C-box, while the Type II promoter consists of an A-box and a B-box ^6^. Generally, Pol III transcription initiation requires transcription factors (TFs), including TFIIIC and TFIIIB ^7^. TFIIIA is a specific TF for Type I promoters and consists of nine zinc-finger (ZF) repeats ^8^. It is the first factor that recognizes and binds the 5S rRNA promoter. The large, six-subunit TFIIIC is recruited to the Type I promoter via TFIIIA. In the case of type II promoters, TFIIIC can directly recognize and bind to A-box and B-box elements, and recruits TFIIIB, positioning it upstream of the transcription start site (TSS) ^9^. TFIIIB, which consists of three subunits – TATA-box binding protein (TBP), TFIIB-related factor (Brf1), and B double prime factor (B”) – may support Pol III transcription alone once stably assembled on the promoter ^9^. The six subunits of the TFIIIC complex are organized into two lobes: subunits τ131, τ95, and τ55 form the τA lobe, and subunits τ138, τ91, and τ60 form the τB lobe. The two lobes are proposed to be connected via a flexible linker that helps TFIIIC bind ICRs of different lengths ^10,11^. While several structures of TFIIIC subcomplexes and domains are solved, including the τA lobe ^12^, the τ131 N-terminal tetra-trico peptide repeats (TPR) array ^10^, the histidine phosphatase domain (HPD) of τ55 ^13^, the τ138 extended winged-helix (eWH) domain ^10^, and a subcomplex of τ60 and τ91 ^14^, the structure of the complete TFIIIC complex remains elusive, possibly due to its high flexibility. Structures of TFIIIB components have been solved as a part of the Pol III transcription pre-initiation complex (PIC) ^15-17^. To date, structures of TFIIIA include ZF 1-3 bound to DNA ^18^, ZF 4-6 bound to 5S rRNA ^19^, and ZF 1-6 bound to 5S rRNA gene ^20^. However, the full-length structure of TFIIIA has not been solved.

Misregulation of Pol III transcription has been linked to cancer ^21-24^, with changes in the expression of TFIIIC subunits being associated with infection and disease ^25^. Several TFIIIC subunits have been found to be overexpressed in ovarian tumors ^26^, and TFIIIC’s occupancy at tDNAs has been shown to increase under stress conditions ^27^. Additionally, research suggests that TFIIIC bound to extra TFIIIC (ETC) sites may play a role in chromosome organization ^28^. Despite the importance of these findings, the mechanism by which TFIIIC recruits Pol III to its promoters is not well understood.

Here, we have used cryo-electron microscopy (cryo-EM) to visualize an *S. cerevisiae* complex of TFIIIA and TFIIIC bound to the 5S rRNA gene in the absence of TFIIIB. Additionally, we present a cryo-EM structure of TFIIIA, TFIIIC, and TFIIIB subunits, Brf1 and TBP, bound to the 5S rRNA promoter. We were able to identify all nine TFIIIA ZFs and locate all six subunits of TFIIIC within the complex. Our structure demonstrates how the two largest subunits, τ138 and τ131, hold together the two lobes of TFIIIC. The full-length 5S rRNA gene, which is 151 base pairs (bp), is wrapped around the TFIIIA-TFIIIC complex. Brf1-TBP binds the upstream region of DNA and contacts the N-terminal region of τ131. We also verify our structural model using single-molecule Förster resonance energy transfer (smFRET) assay, which reveals the dynamic nature of the DNA bound to the complex. We discuss the role of TFIIIC in Pol III PIC assembly and propose a model for how the TFIIIA-TFIIIC complex, bound to ICR, may help TFIIIB in finding its binding site upstream of the TSS.

## RESULTS

### Brf1-TBP stabilizes the TFIIIC-TFIIIA complex

The TFIIIA-TFIIIC and TFIIIA-TFIIIC-Brf1-TBP complexes were assembled on a double-stranded (ds) DNA template composed of the *S. cerevisiae* 5S rRNA gene, including the upstream TFIIIB-binding site and the gene body containing ICR (Figure 1A, see Methods). The complexes were assembled in a step-wise manner using individually purified *S. cerevisiae* factors. Cryo-EM datasets were collected for both complexes, with a subset of 109,548 particles refined to 6.6 Å resolution for the TFIIIA-TFIIIC complex (Figure 1B; Figure S1), and a subset of 78,512 particles refined to 3.8 Å resolution for the TFIIIA-TFIIIC-Brf1-TBP complex (Figure 1C; Figure S2; Movies S1 and S2).

**Figure 1.**
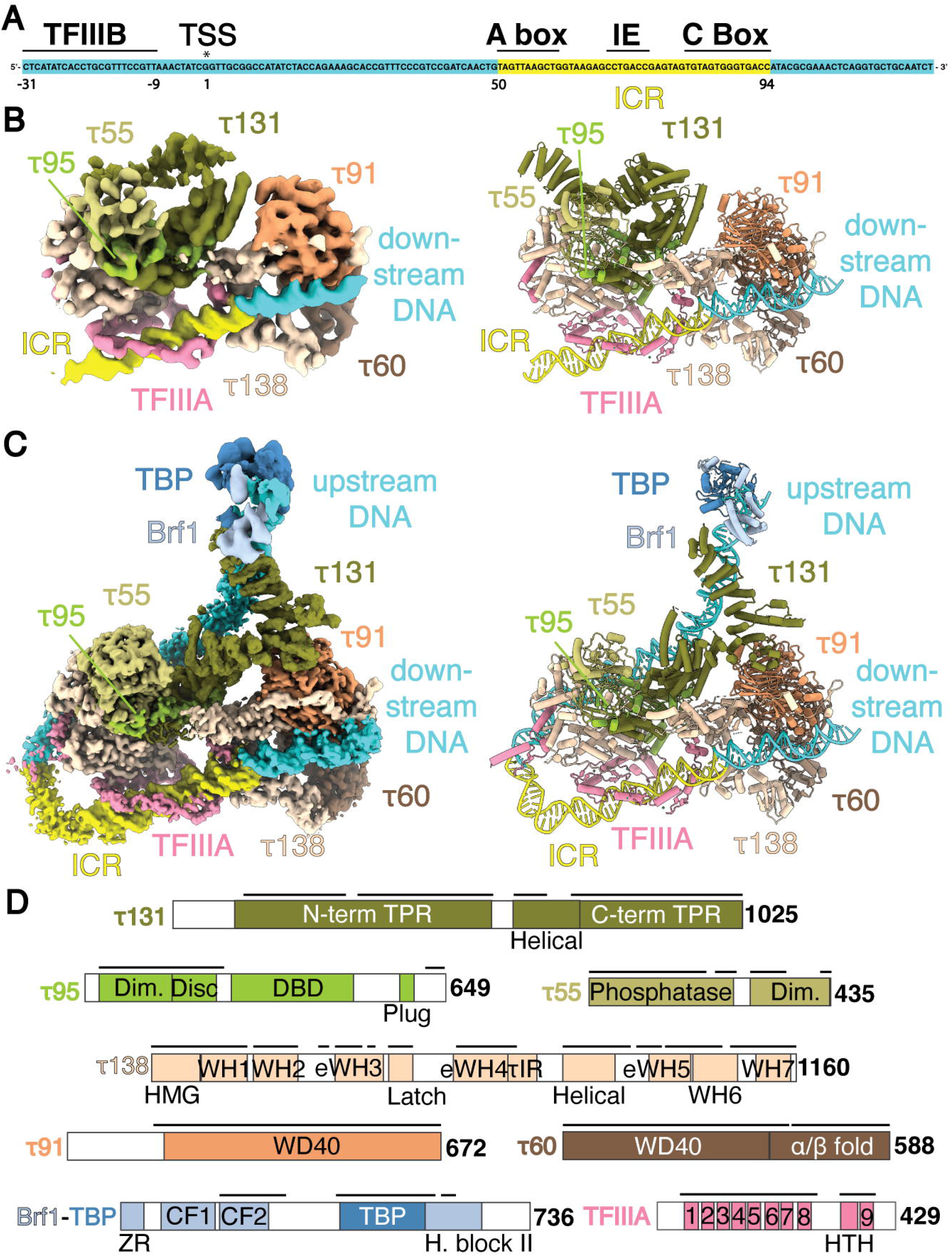
Cryo-EM structures of the TFIIIA-TFIIIC complex and the TFIIIA-TFIIIC-Brf1-TBP complex bound to the 5S rRNA gene. (A) Schematic representation of the DNA template, with transcription start site (TSS) and intermediate element (IE) indicated. (B) Density map of the TFIIIA-TFIIIC complex (left) and the corresponding model of the TFIIIA-TFIIIC complex (right). The internal control region (ICR) is highlighted. (C) Composite density map of the TFIIIA-TFIIIC-Brf1-TBP complex (left, see Methods) and the corresponding model of the TFIIIA-TFIIIC-Brf1-TBP complex (right). (D) Schematic domain representation of TFIIIA, TFIIIC, and Brf1-TBP subunits. The amino-acid lengths of the subunits are labeled at the C-termini. Black lines above the bars show the portions for which models were built. Dim., dimerization domains; DBD, DNA-binding domain; HMG, high mobility group box domain; WH, winged helix; eWH, extended winged helix; τIR, τ131-interaction region; ZR, zinc ribbon; CF1,2, cyclin fold 1,2; H. block II, homology block II; TFIIIA 1-9, TFIIIA zinc fingers 1-9; HTH, helix-turn-helix. See also Figures S1, S2, S3; Table 1; Movies S1 and S2.

The TFIIIA-TFIIIC and TFIIIA-TFIIIC-Brf1-TBP complexes are assembled on identical DNA templates. The overall conformation of TFIIIC is similar between the two complexes, and the ZF array of TFIIIA interacts with the ICR in a similar manner (Figure 1, B and C; Figure S3A). All six subunits of TFIIIC are visible in both complexes (Figure 1, B to D). The two lobes of TFIIIC, τA and τB, are in close contact with each other, held together by multiple interactions with DNA and the τ138 subunit, which is shared between the two lobes. The τA lobe is comprised of a τ95-τ55 dimer, the C-terminal TPR array of τ131, and the C-terminal half of τ138. The τB lobe includes subunits τ91, τ60 and the N-terminal half of τ138. The interaction between τA and TFIIIA, as well as those between τB and DNA, are identical in the two complexes.

**Table 1.**
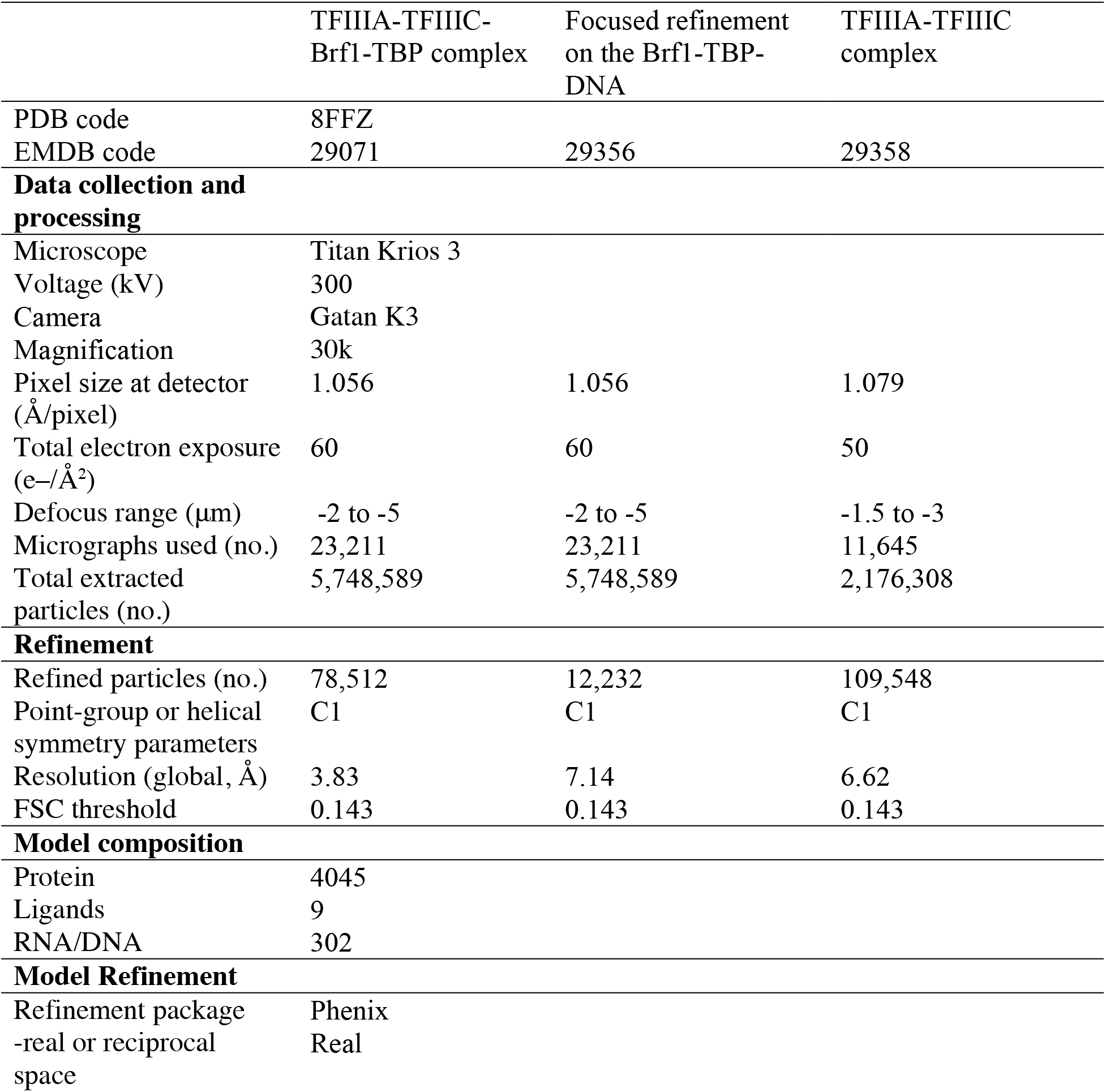
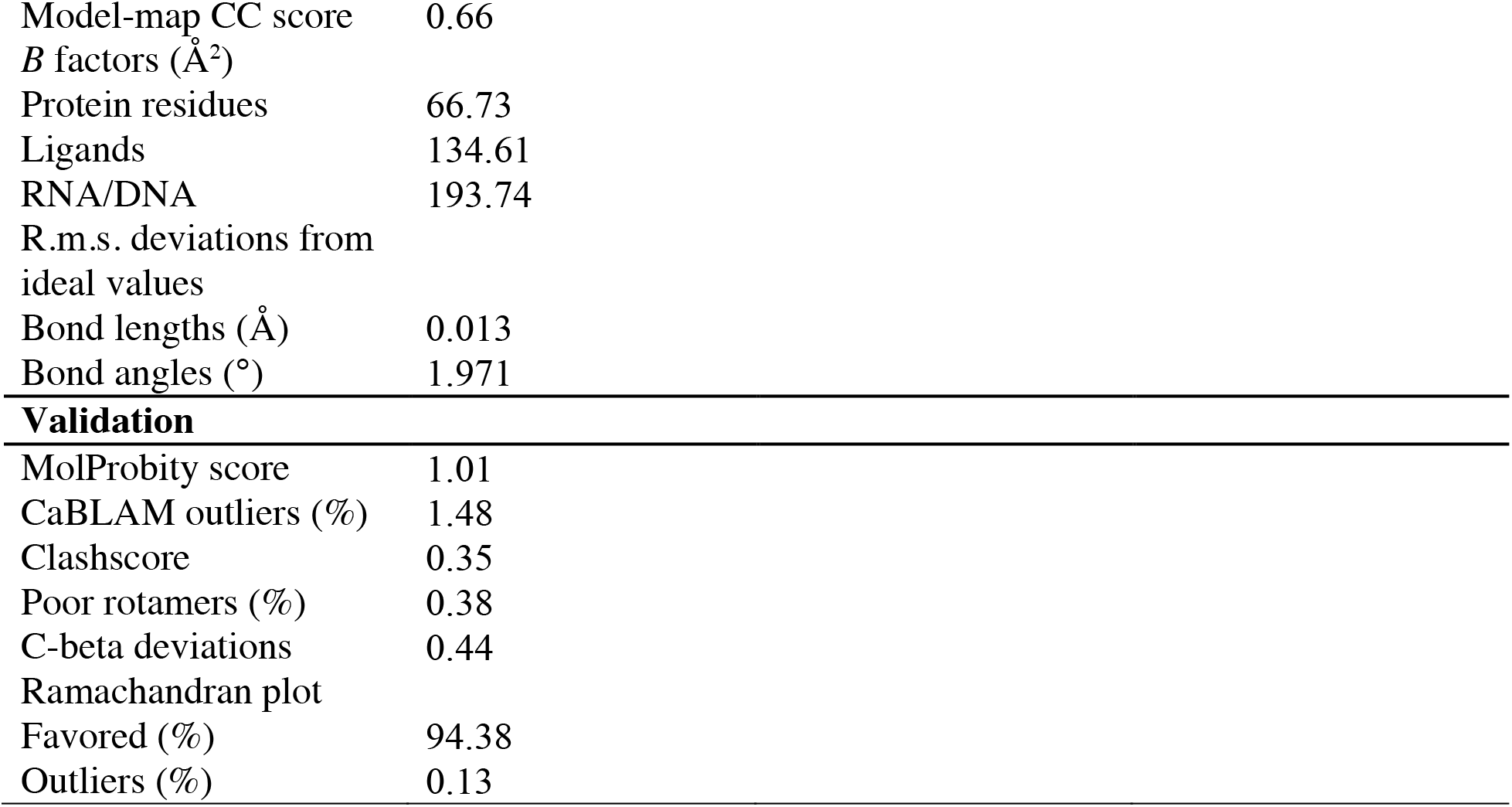
Cryo-EM data collection, refinement, and validation statistics.

|  | TFIIIA-TFIIIC-Brf1-TBP complex | Focused refinement<br>on the Brf1-TBP-DNA | TFIIIA-TFIIIC<br>complex |
| --- | --- | --- | --- |
| PDB code | 8FFZ |  |  |
| EMDB code | 29071 | 29356 | 29358 |
| **Data collection and processing** |  |  |  |
| Microscope | Titan Krios 3 |  |  |
| Voltage (kV) | 300 |  |  |
| Camera | Gatan K3 |  |  |
| Magnification | 30k |  |  |
| Pixel size at detector<br>(Å/pixel) | 1.056 | 1.056 | 1.079 |
| Total electron exposure<br>(e-/Å<sup>2</sup>) | 60 | 60 | 50 |
| Defocus range (μm) | -2 to -5 | -2 to -5 | -1.5 to -3 |
| Micrographs used (no.) | 23,211 | 23,211 | 11,645 |
| Total extracted<br>particles (no.) | 5,748,589 | 5,748,589 | 2,176,308 |
| **Refinement** |  |  |  |
| Refined particles (no.) | 78,512 | 12,232 | 109,548 |
| Point-group or helical<br>symmetry parameters | C1 | C1 | C1 |
| Resolution (global, Å) | 3.83 | 7.14 | 6.62 |
| FSC threshold | 0.143 | 0.143 | 0.143 |
| **Model composition** |  |  |  |
| Protein | 4045 |  |  |
| Ligands | 9 |  |  |
| RNA/DNA | 302 |  |  |
| **Model Refinement** |  |  |  |
| Refinement package | Phenix |  |  |
| -real or reciprocal<br>space | Real |  |  |

|  |  |
| --- | --- |
| Model-map CC score | 0.66 |
| *B* factors (Å<sup>2</sup>) |  |
| Protein residues | 66.73 |
| Ligands | 134.61 |
| RNA/DNA | 193.74 |
| R.m.s. deviations from ideal values |  |
| Bond lengths (Å) | 0.013 |
| Bond angles (°) | 1.971 |
| **Validation** |  |
| MolProbity score | 1.01 |
| CaBLAM outliers (%) | 1.48 |
| Clashscore | 0.35 |
| Poor rotamers (%) | 0.38 |
| C-beta deviations | 0.44 |
| Ramachandran plot |  |
| Favored (%) | 94.38 |
| Outliers (%) | 0.13 |

The presence of Brf1-TBP in the complex alters the interactions between TFIIIC τA lobe and DNA (Figure 1, B and C). In the TFIIIA-TFIIIC-Brf1-TBP complex, the DNA upstream of ICR is visible and is stabilized by multiple interactions with TFIIIC τA lobe. The addition of Brf1-TBP dramatically changes the position and stability of τ131 N-terminal TPR array. This part of τ131 is not well-resolved in the TFIIIA-TFIIIC complex, possibly due to its unconstrained movement relative to the rest of the complex (Figure S3, C and D). The residues connecting the N-terminal and C-terminal TPR arrays are not resolved in both structures, suggesting that the two arrays are connected by a flexible ‘hinge’ domain. Interactions between τ131 N-terminal TPR, upstream DNA, and Brf1-TBP lead to the extended and stabilized conformation of τ131. This state is additionally stabilized by contacts between the ‘ring’ domain of τ131 (residues 390 - 428) and τ91 (Figure 1, B and C). The well-positioned upstream DNA leads to the stabilization of TFIIIA ZF 9. The TFIIIA-TFIIIC-Brf1-TBP complex is resolved to higher resolution, thus it is used to describe protein-protein interactions within TFIIIC.

### τ138 bridges TFIIIC τA and τB lobes

The two largest subunits of TFIIIC, τ138 and τ131, facilitate interactions between the τA and τB lobes (Figure 2, A and B). This is consistent with previous genetic study that identified functional connections between τ138 and τ131 ^29^. These two large subunits also play a role in supporting interactions between subunits within each lobe.

**Figure 2.**
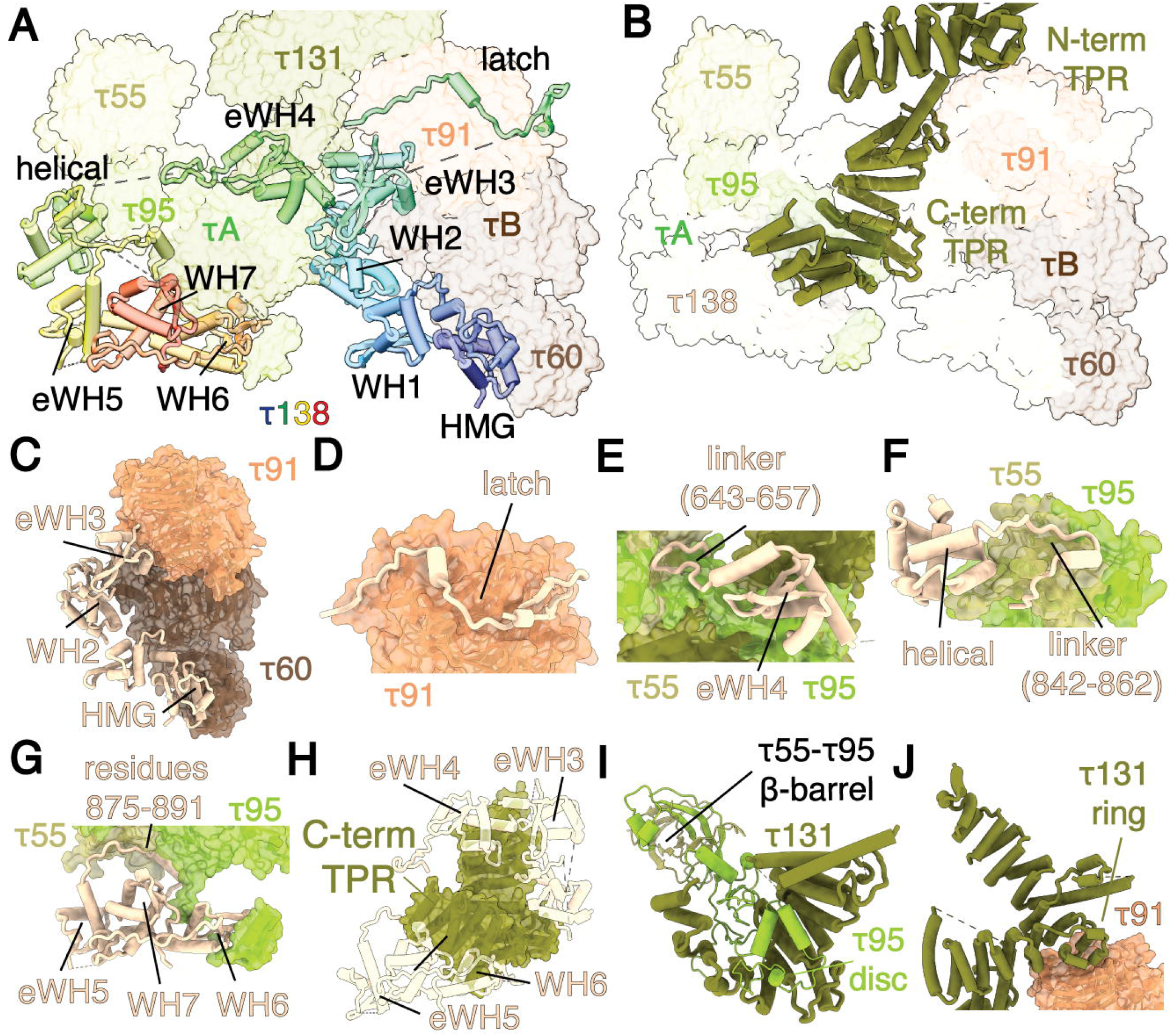
Bridging of TFIIIC τA and τB lobes by the two largest subunits, τ138 and τ131. (A) The τ138 subunit acts as a scaffold for the body of TFIIIC and mediates the interaction between τA and τB lobes. The τ138 subunit is shown as cartoon, while other TFIIIC subunits are shown as transparent surfaces. The τ138 subunit is colored from N-terminus to C-terminus in a rainbow pattern. (B) The τ131 subunit is located in the middle of the complex. The C-terminal TPR array of τ131 is buried in the body of the complex, while the N-terminal TPR array is extended away from the body of the TFIIIA-TFIIIC-Brf1-TBP complex. The τ131 subunit is shown as cartoon, with other TFIIIC subunits depicted as transparent surfaces. (C) The WD40 domain of τ60 supports the WH2 and eWH3 domains of τ138, while the α/β fold of τ60 contacts the HMG domain of τ138. The WD40 domain of τ91 supports the eWH3 domain of τ138. (D) The latch of τ138 (residues 425-470) is attached to the surface of τ91. (E) The eWH4 domain of τ138 contacts the helical domain of τ131, followed by a linker (residues 643-657) binding to the τ55-τ95 dimer. (F) The helical domain and linker (residues 842-862) of τ138 bind to τ55. (G) The residues 875-891 of τ138 contact the τ55-τ95 dimer. The eWH5, WH6 and WH7 domains form a compact fold, interacting with the τ55-τ95 dimer and the C-terminal region of τ95 (residues 612-647). (H) Contacts between τ138 and τ131. The C-terminal TPR of τ131 interacts with eWH4 (residues 1000-1023) and WH7 of τ138. (I) Interactions within τA lobe: the dimerization domains of τ55 and τ95 subunits form a β-barrel. The disc domain of τ95 interacts with the C-terminal TPR array of τ131. (J) The N-terminal TPR array of τ131 is in the extended state, protruding away from the C-terminal TPR array and the body of TFIIIC. This position is stabilized through interactions of the τ131 ring domain with the τ91 subunit.

The domains of τ138, the largest TFIIIC subunit, are distributed between the two lobes (Figure 2A). The N-terminus of τ138 belongs to the τB lobe, the middle region is situated in the center, and C-terminus is a part of the τA lobe. The compact τA and τB regions of τ138 are connected by a less structured region (residues 418-739) that contains the eWH4 domain in the middle. Part of this region (residues 641-693) has been shown to be the main link between the two lobes of TFIIIC ^10^. Subunit τ138 comprises seven WH domains, three of which are eWH (Figures 1D, 2A). The HMG, WH2, and eWH3 domains bind to τ60 subunit, and eWH3 contacts τ91 (Figure 2C), consistent with previous biochemical work ^14,30^. The ‘latch’ domain of τ138 (residues 425-470) is attached to the surface of τ91 WD40 (Figure 2D). The eWH4 domain interacts with the helical domain of τ131 (residues 612 - 732), and the linker (residues 643-657) connects eWH4 with the τ55-τ95 dimer (Figure 2E). The helical domain of τ138 resides in the τA lobe and interacts with the τ55-τ95 dimer (Figure 2F). eWH5, WH6, and WH7 form a compact structure, interacting with the τ55-τ95 dimer and C-terminal region of τ95 (residues 612-647) (Figure 2G), consistent with previous crosslinking mass spectrometry results ^10^. Subunit τ138 serves as hub that brings together all other parts of the complex by interacting with five other TFIIIC subunits, TFIIIA and DNA (Figures 2A, 3C, 4B). Additionally, the two largest subunits, τ138 and τ131, also interact with each other (Figure 2H).

τ131 is a component of the τA lobe and is located in the middle of the structure (Figure 2B). The TPRs of τ131 are divided into two modules: N-terminal TPR and C-terminal TPR. The N-terminal TPR is further subdivided into two ‘arms’ with a ‘ring’ domain between them ^10,31^.

The concave surface of the C-terminal τ131 TPR array accommodates the ‘disc’ domain (residues 161-236) of τ95 (Figure 2I), which is in agreement with previous studies in yeast and humans ^10,12,32^. The ring domain of N-terminal TPR also contacts τ91 (residues 259-280) (Figure 2J). Notably, the area of contact between the lobes is smaller in comparison to the area of contact within each lobe (Figure 2, C to J). τB lobe has a buried surface area of 6669 Å^2^, while τA lobe buries a large area of 15828 Å^2^. However, the buried surface area between τA and τB is only 675 Å^2^.

### 5S rRNA gene wraps around the complex

The DNA construct used in the experiment is composed of the 5S rRNA gene sequence extended to the upstream TFIIIB-binding region (Figure 1A) ^33,34^. The TFIIIB-binding sequence (bp -31 to -9) is followed by the TSS (position 1), the ICR (bp 50 to 94), and the downstream region. The location of Brf1-TBP and TFIIIA are in excellent agreement with the registers of the TFIIIB-binding site and the ICR in the 5S rRNA, as determined by tracing the DNA density in the full complex reconstruction. The DNA wraps around the complex and makes multiple contacts with TFIIIA, TFIIIC, and Brf1-TBP (Figure 3; Figure 4A). The path of the bound DNA appears to make a roughly 180 degrees turn, bringing the upstream and downstream regions closer together. The ICR is the most sharply bent region of the DNA (Figure 3A). These findings align with previous research in which both TFIIIA and TFIIIC were shown to introduce bending within DNA ^35-38^.

**Figure 3.**
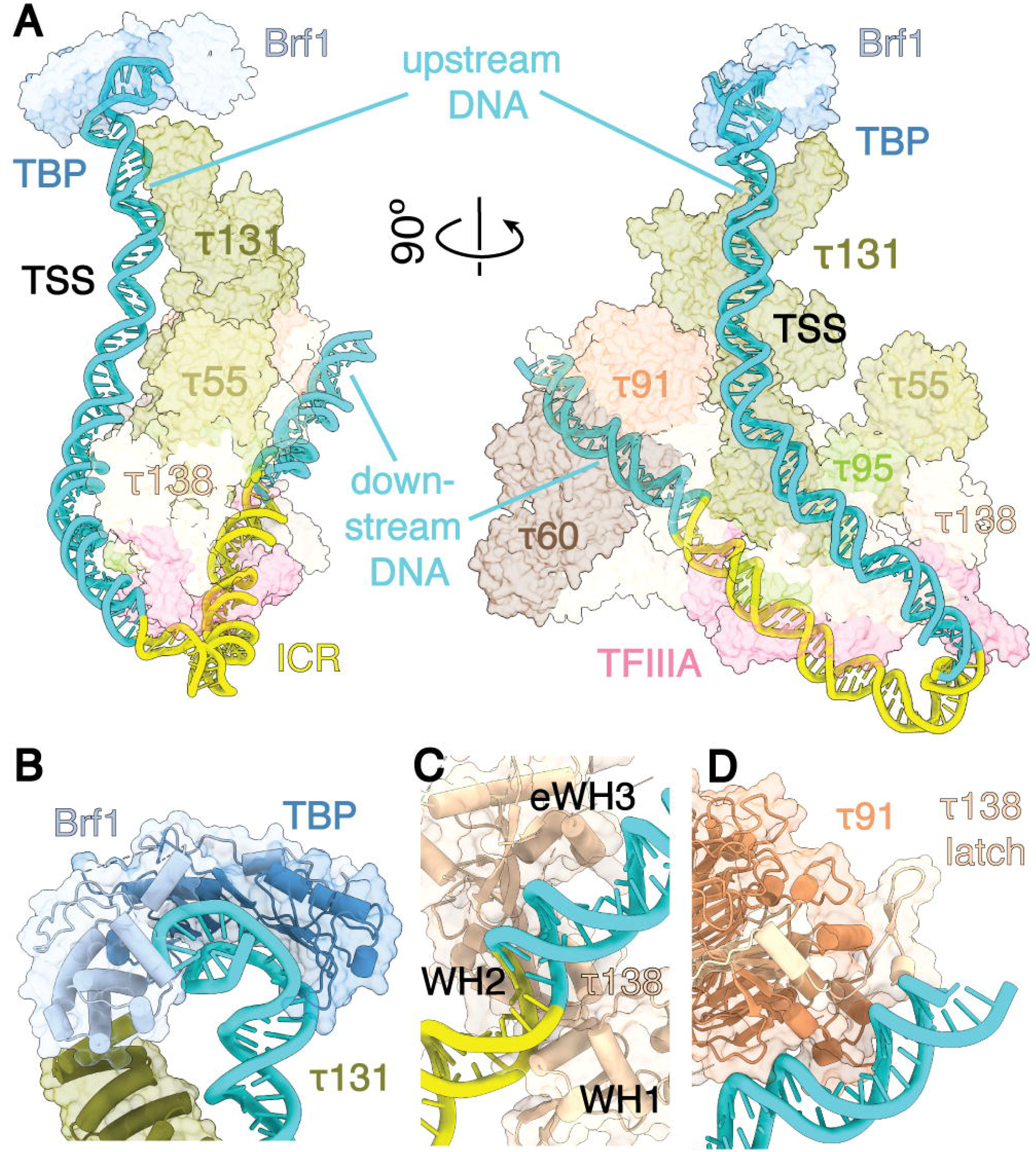
Brf1-TBP binding leads to the 5S rRNA gene wrapping around the TFIIIA-TFIIIC body. (A) DNA wraps around the TFIIIA-TFIIIC-Brf1-TBP complex. The largest protein surfaces that interact with DNA are τ131 and TFIIIA. DNA is shown as cartoon, and proteins are shown as transparent surfaces. (B) TBP bends upstream DNA region. This part of DNA is stabilized by τ131, TBP, and Brf1. (C) τ138 WH1, WH2, and eWH3 wrap around downstream DNA. (D) The end of the 5S rRNA gene is supported by the τB lobe. Both τ91 and the τ138 latch (residues 425-470) are associated with this part of the DNA.

**Figure 4.**
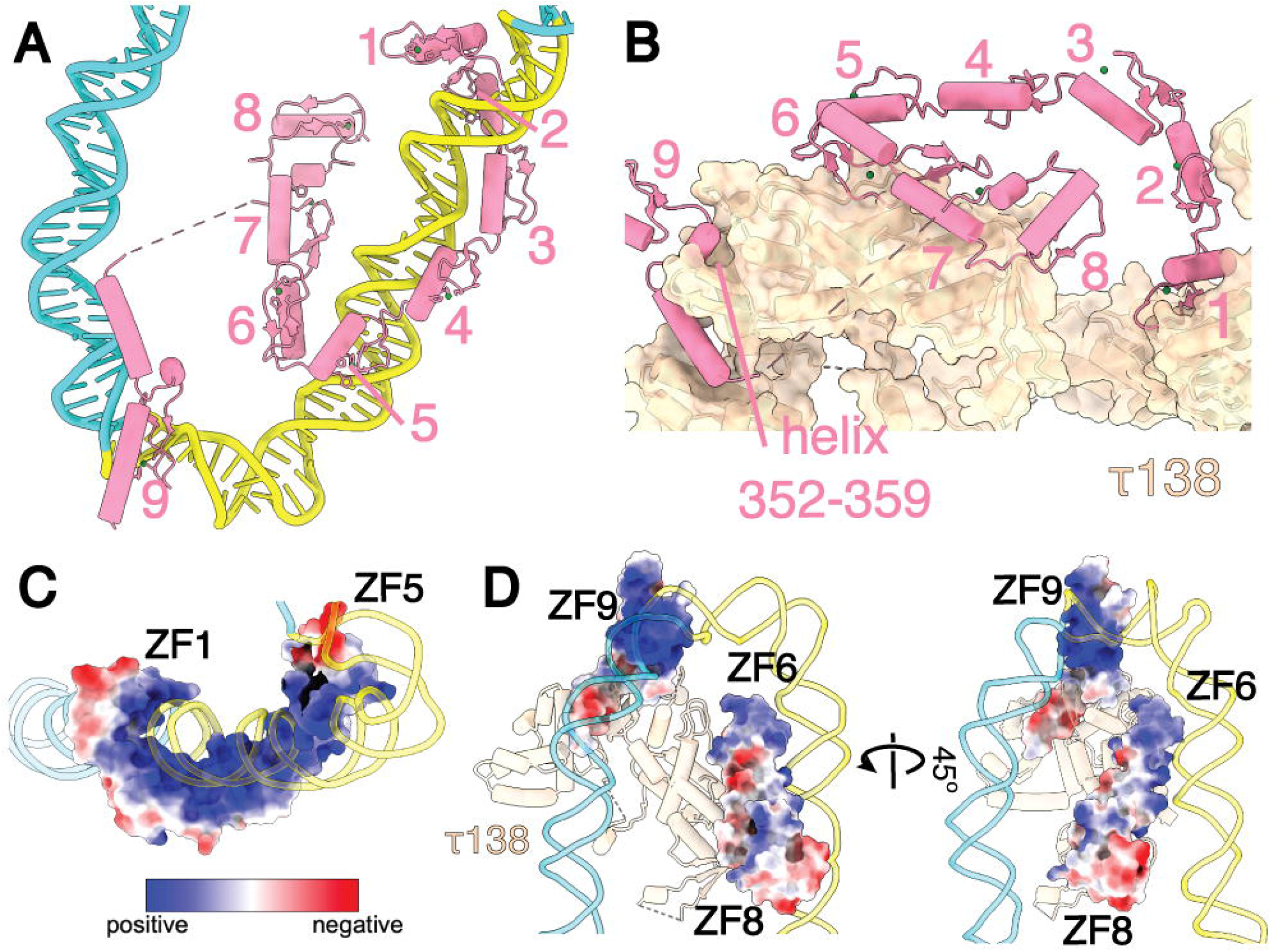
TFIIIA binds the ICR of the 5S rRNA gene. (A) ZF 1 to 5 and ZF 9 bind DNA in the ICR, while ZF 6-8 protrude away from ICR. The most sharply bent region of DNA is located between ZF 5 and 9. (B) TFIIIA has a large interaction surface with τ138. ZF 1, 6-8, and a helix near ZF 9 contribute to the interaction. (C) DNA-binding residues of ZF 1 to 5 form a large, extended, positively charged surface. TFIIIA is shown as surface and colored according to coulombic potential. The model is colored in a range from red for negative potential to blue for positive potential. (D) ZF 6 to 8 do not interact with DNA as closely as the first five ZFs, but their positively charged surfaces are directed towards DNA and away from τ138 (E). ZF9 DNA binding surface is positively charged. TFIIIA is shown as surface and colored according to coulombic potential. The model is colored in a range from red for negative potential to blue for positive potential. See also Figure S5.

The upstream TFIIIB-binding region is recognized by TBP and Brf1 ^34,39^. The resolution of this part of the map is not sufficient to allow *de novo* model building, so the Brf1-TBP-DNA model (PDB ID 6cnb) was docked in the map as rigid body (Figure 3B; Figure S3B). The DNA upstream of the ICR is bound to positively charged patches on the τA lobe, primarily τ131 (residues 192 to 252; 667 to 718; 832 to 841; 924 to 931) (Figure 3A; Figure S4, A and B). The binding of τ131 to the upstream region of the *S. cerevisiae* 5S rRNA and SUP4 tRNA Tyr genes has also been previously shown through site-specific DNA-protein photocrosslinking ^39,40^. The first three WH domains of τ138 are wrapped around the DNA downstream of the ICR, following the minor groove of DNA (Figure 3C). The positively charged regions of WH1 and eWH3 have close contacts with the DNA minor and major groove, and similar examples of WH-DNA interactions can be found in other transcription initiation complexes (Figure S4D). The DNA-binding surfaces of τ138 and τ131 have highly conserved residues forming positively charged patches (Figure S4, A to C; Figure S4, E and F). The downstream region of the DNA is bound by the WD40 domain of τ91 subunit (Figure 3D; Figure S4C). This subunit has been shown to photo-crosslink to the very end of the 5S rRNA gene ^39^, while the *S. pombe* homolog of τ91, Sfc6p, has been shown to recognize the B-box in Type II promoter ^41^. This part of the DNA is additionally supported by the τ138 latch (residues 449 - 470) (Figure 3D).

### TFIIIA binds the ICR of the 5S rRNA gene

DNA footprinting assay has revealed that both *X. laevis* and *S. cerevisiae* TFIIIA bind to the ICR of the 5S rRNA gene ^34^. DNA binding of the first three ZF in our model is identical to the *X. laevis* TFIIIA-DNA structures but deviates for the following ZF, likely due to the presence of its binding partner, TFIIIC, in the full complex (Figure S5A) ^20,34,42^. All nine ZFs share the same fold, but ZF 9 has a longer helix (Figure S5B). The ICR is protected by TFIIIA ZF 1 to 5 and ZF 9, while ZF 6 to 8 point away from DNA (Figure 3A; Figure 4A). Previous study has shown that purified TFIIIA protects the ICR of the 5S rRNA gene (bp 66 to 95) from DNase I cleavage, with enhanced cleavage at bp 50 and 65, consistent with our structural model ^34^. The footprints of the five N-terminal ZF bound to 5S rRNA gene were indistinguishable from binding of full-length TFIIIA, and it was previously suggested that ZF 6 to 9 do not bind DNA tightly ^42-44^. Consistently, ZF 1 to 3 in our model bind in the DNA major groove (Figure 4A; Figure S5C) ^18,45^. ZF 4 traverses the minor groove, while ZF 5 and 9 bind the major groove again (Figure 4A; Figure S5C). The sharpest part of the DNA bend is located between ZF 5 and ZF 9 (Figure 4A). All DNA-binding ZF show positive charge and high conservation of their DNA-contacting surface (Figure 4, C and D; Figure S5D).

τ138 was previously suggested to bracket TFIIIA on 5S rRNA gene ^39^. Our observation reveals that the contact between TFIIIA and TFIIIC is maintained through τ138, with a large surface area between ZF 6-8 of TFIIIA and residues 980-1072 of τ138 (Figure 4B). Additionally, ZF 1 and helix-turn-helix domain (residues 331 - 363) of TFIIIA also contribute to this interaction. This region within the C-terminus of *Xenopus* TFIIIA has been identified as a TFIIIC binding and non-DNA binding site ^46^. ZF 7 was found to be essential for the assembly of transcriptionally active complex ^47^. The presence of ZF 7 to 9 has also been shown to be necessary for the transcription activity of the complex, likely due to higher-order interactions in the complex ^43,44^. A flexible linker between ZF 8 and 9 is not visible in the map, except for the helix-turn-helix domain (residues 331 - 363) right next to ZF 9 (Figure 4A). TFIIIA lacking this region (residues 283 - 364) has been shown to be able to recruit TFIIIC but unable to promote transcription ^43^. Specifically, the deletion of a leucine-rich segment 352-NGLNLLLN-359, a small helix next to ZF 9, resulted in the loss of transcription activity ^48^. This helix appears to be an anchor point of ZF 9 on the surface of τ138 (Figure 4B).

### smFRET shows dynamic nature of the complex

To understand the conformational dynamics of the complex and independently verify our structural model, we perform smFRET assay using the full TFIIIA-TFIIIC-Brf1-TBP complex (Figure 5A). We purified the complex using a dsDNA molecule labeled with Cy3 and Cy5, following the same protocol as for the cryo-EM sample preparation (see Methods). The positions of Cy3 and Cy5 were chosen such that, based on our structure, they are 66 Å apart in the assembled complex, which would allow for efficient FRET (Figure 3A). The DNA-only sample shows stable FRET close to zero, indicating an extended conformation of the DNA (Figure 5B). The TFIIIA-TFIIIC-Brf1-TBP complex assembled on the DNA shows two higher FRET states (Figure 5C). The FRET histogram shifts to higher FRET, indicating a decrease in the distance between the fluorophores (Figure 5D). Notably, most traces (62%) display slow transitions between the two FRET states at 0.07 and 0.22 before photobleaching (Figure 5, C and E; Figure S6). The second FRET peak for the full complex at a FRET value of 0.22 corresponds to a generic distance of 66 Å, which is consistent with our prediction from the structure, confirming that this is the wrapped DNA conformation. To further investigate the nature of the low FRET state, we conduct the experiment on the complex that is assembled without Brf1-TBP. The FRET histogram for this complex shows a single major peak at a FRET value of 0.07, similar to the lower FRET peak for the full complex, suggesting that this represents a dynamic, partially unwrapped intermediate (Figure 5D). The presence of this lower FRET state in TFIIIA-TFIIIC-Brf1-TBP complex trajectories suggests that this complex is dynamic and the upstream DNA is not always bound by both TFIIIC and Brf1-TBP.

**Figure 5.**
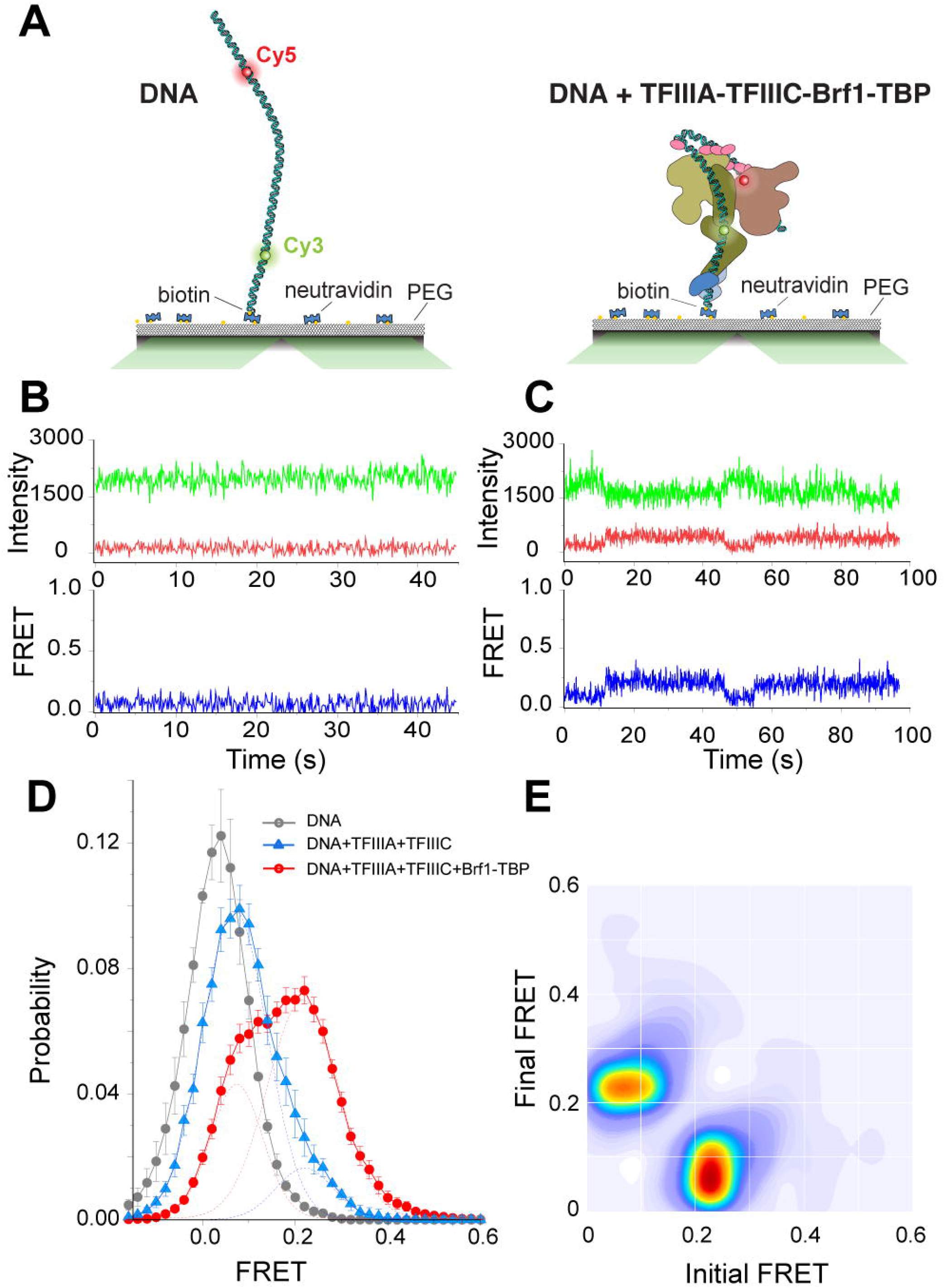
smFRET reveals DNA bending that is consistent with the cryo-EM study. (A) Schematic representation of the smFRET assay. Left, labeled DNA; right, labeled DNA + TFIIIA-TFIIIC-Brf1-TBP complex. (B) Representative single-molecule time traces of DNA only showing donor (green) and acceptor (red) intensities and the corresponding FRET (blue). (C) Representative single-molecule time traces of the TFIIIA-TFIIIC-Brf1-TBP complex showing donor (green) and acceptor (red) intensities and the corresponding FRET (blue). The FRET data shows the presence of two states. (D) smFRET population histogram of DNA only, DNA + TFIIIA-TFIIIC, and DNA + TFIIIA-TFIIIC-Brf1-TBP. Data represents mean±s.e.m. The histogram of DNA + TFIIIA-TFIIIC is fitted to two Gaussian distributions (blue dashed line) centered on 0.07 and 0.22, respectively. The histogram of the full complex, DNA + TFIIIA-TFIIIC-Brf1-TBP, is also fitted to two Gaussian distributions (red dashed line) centered on 0.07 and 0.22, respectively. (E) Transition density plot of the full complex, DNA + TFIIIA-TFIIIC-Brf1-TBP. Transitions are from two independent measurements. See also Figure S6.

### TFIIIC-mediated assembly of Pol III PIC

Based on the cryo-EM structures and smFRET data, we propose a model for the TFIIIC-mediated assembly of the Pol III PIC. Before the complex assembles, the two lobes of TFIIIC, connected by the τ138 linker, may move relative to each other (Figure 6, state 1). This relative flexibility of the lobes have been previously observed by EM ^11^. We also observe different positions of the TFIIIC lobes by negative stain EM (Figure S7A). The assembly of the Pol III PIC is thought to begin with TFIIIA locating the 5S rRNA gene ^34,49,50^, likely introducing initial bending that is mostly localized within the ICR ^36,37^. In our structure, only ZF 1 to 5 and ZF 9 are tightly bound to DNA. However, when TFIIIA first binds DNA, all nine ZF may make DNA contacts, similar to *X. laevis* TFIIIA ^51,52^. In our structure, ZF 1 to 5 bind 29 bp, while ZF 9 protects 5 bp of the ICR. This leaves about one turn of DNA (11 bp) that, in principle, can be occupied by ZF 6 to 8 in this initial phase. The binding of ZF 6 to 8 may be similar to the DNA-binding mode of ZF 1 to 3. Compared to the first five ZF, ZF 6 to 8 have smaller positively charged areas, which may make them easier to be peeled off from DNA when the TFIIIA-TFIIIC complex forms (Figure 4, C and D). This first ‘searching’ state can be characterized by the flexibility of the TFIIIA-DNA complex and within the unrestrained TFIIIC.

**Figure 6.**
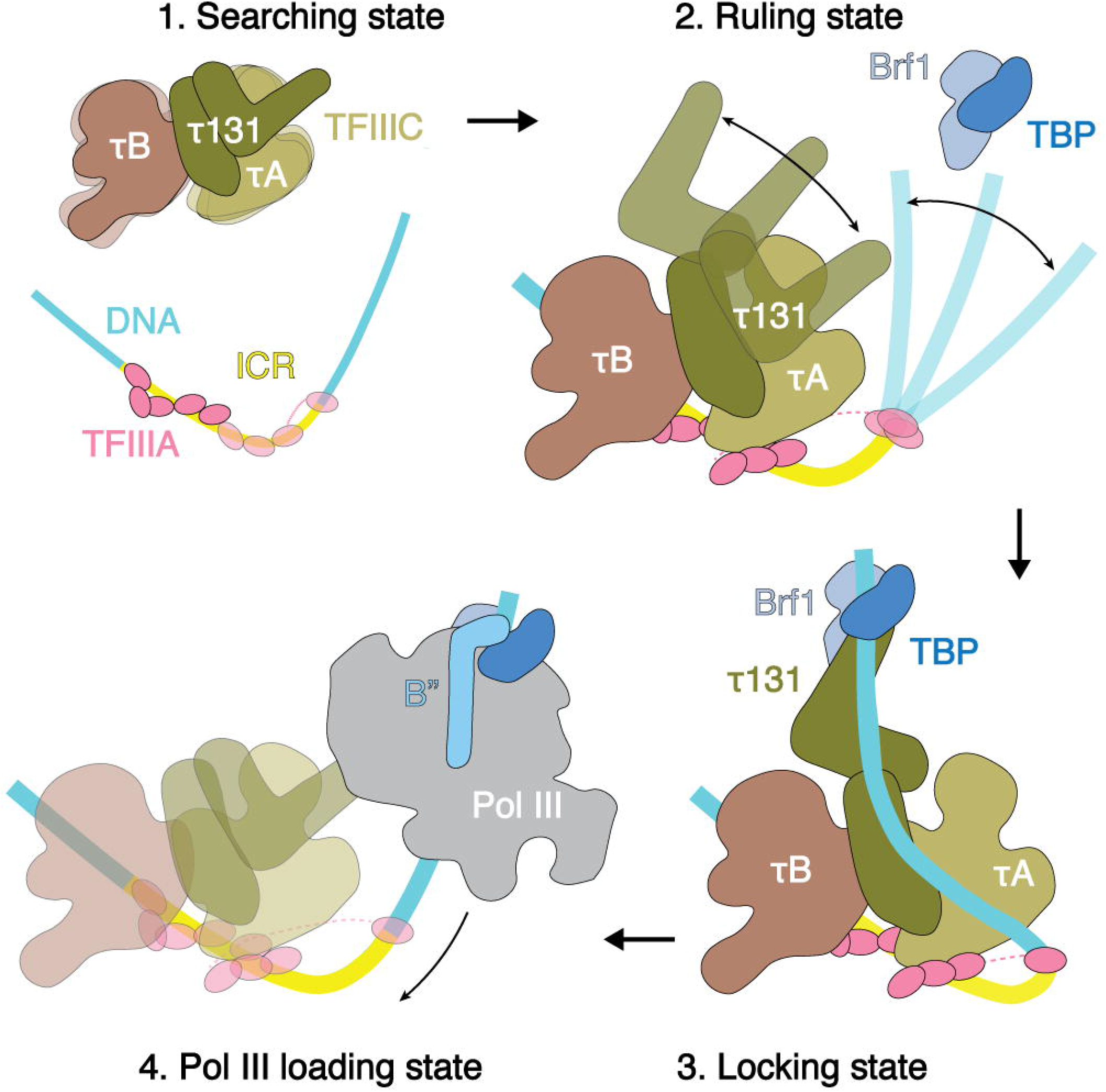
Model of TFIIIC-dependent Pol III PIC assembly on 5S rRNA gene. 1. Searching state: The two lobes of TFIIIC are not restrained by DNA binding and may move relative to each other. TFIIIA recognizes the ICR of the 5S rRNA gene, and this binding can introduce initial bending in the DNA, localized within ICR. 2. Ruling state (formation of the TFIIIA-TFIIIC-DNA complex): TFIIIC binds TFIIIA and DNA downstream of ICR. These interactions restrain mobility between the τA and τB lobes of TFIIIC and stabilize the body of the complex. The ‘hinge’ region between the N-terminal TPR and C-terminal TPR of τ131 allows the N-terminal TPR to transition from the closed state (N-terminal TPR contacts C-terminal TPR) to the fully open state (N-terminal TPR array is turned by 180 degrees relative to its position in the closed state). This sampling movement may help τ131 N-terminal TPR to search for Brf1 and/or DNA for binding. The ICR and downstream DNA is fixed via interactions with ZF 1-5 of TFIIIA and the τB lobe. The flexibility within the upstream DNA helps to search for τ131 and/or Brf1 binding. This complex may represent the ‘ruling’ state of τ131 because the distance between the N-terminal and C-terminal TPRs of τ131 is variable, but strong τ131-DNA interaction is only possible when N-terminal TPR is in the extended conformation. This allows τ131 to measure the distance from ICR to the TFIIIB-binding region. 3. Locking state (formation of the TFIIIA-TFIIIC-Brf1-TBP-DNA complex): N-terminal TPR array of τ131 is in the extended state. DNA is bent within the ICR, and upstream DNA is bound to the surface of the τA lobe. DNA upstream of TSS is bound by TBP, Brf1, and the N-terminus of τ131. 4 Pol III loading state: Once Brf1 and TBP locate the TFIIIB-binding site and bind it, B” and Pol III can be recruited. Pol III may then initiate transcription, while TFIIIC and TFIIIA have to be displaced from DNA to allow Pol III to transcribe the full length of the gene. See also Figure S7.

Once TFIIIC binds TFIIIA and downstream DNA, the mobility between the τA and τB lobes becomes limited (Figure 6, state 2). The ICR and downstream DNA is fixed via interactions with ZF 1-5 of TFIIIA and τB lobe. The upstream DNA may still move due to the presence of a flexible linker within TFIIIA. This movement of upstream DNA can assist in the search for τ131 and/or Brf1-TBP binding. The lower FRET state of 0.07 may correspond to this complex (Figure 5D). The C-terminal TPR array of τ131 is locked in the body of the complex, while the N-terminal TPR array is not restrained. The ‘hinge’ region between the N-terminal and C-terminal TPRs allows for the potential movement of the N-terminal TPR array from the closed state, where it contacts the C-terminal TPR ^12^, to the fully open state, where it is rotated by approximately 180 degrees relative to its position in the closed state (Figure S3D). In this ‘ruling’ phase, the conformational sampling of the τ131 N-terminus may help to search for Brf1-TBP and DNA. The variable distance between the N-terminal and C-terminal TPR arrays of τ131 can act as a ruler: simultaneous interactions of τ131 with TFIIIB and the TFIIIB-binding region of DNA are possible only when τ131 N-terminus is located within a certain distance from the C-terminal TPR (Movie S2). We propose this model of TFIIIC-aided TFIIIB recruitment as an extension of the previously communicated models ^12,53-55^.

The TFIIIA-TFIIIC-Brf1-TBP complex, which we observe after adding Brf1-TBP to TFIIIA-TFIIIC complex, corresponds to ‘locking’ state (Figure 6, state 3). In this state, the N-terminal TPR array of τ131 is fixed in the open conformation. This position of τ131 opens a positively charged patch on its surface (Figure S4, A to C). Yeast two-hybrid assays have shown that Brf1 interacts with the N-terminus of τ131 subunit, and this result has been supported by mutagenesis analysis and binding assays ^31,56-59^. The sharpest bend in DNA is located between TFIIIA ZF 5 and 9 within the ICR region. This bent DNA is stabilized through interactions with TFIIIA and τ131 (Figure 3A; Figure 4A; Figure S4, A to C). Upstream DNA is bound by TBP, Brf1, and the N-terminus of τ131. In the smFRET assay, this state is represented by the 0.22 FRET efficiency state (Figure 5D). DNAse I footprinting has shown that TFIIIA protects the ICR of the 5S rRNA gene, and the binding of TFIIIC to TFIIIA-DNA can extend the footprint in two ways: a ‘core’ footprint on the downstream DNA or ‘extended’ footprint up to upstream DNA region bp -20 ^34^. The addition of TFIIIB extends DNA protection up to bp -45, and the TFIIIA-TFIIIC-TFIIIB complex protects the DNA from bp -45 to bp 120 ^34^. Similar footprinting patterns have been observed for tRNA genes as well ^60^.

The TFIIIA-TFIIIC-Brf1-TBP state that we capture in this study is not compatible with Pol III binding. Alignment of Brf1-TBP from Pol III PIC structures with Brf1-TBP in our model results in severe clashes between Pol III and TFIIIC (Figure S7, B and C). In our model, simultaneous binding of Pol III and TFIIIC to the DNA can only occur if the upstream DNA with Brf1-TBP is detached from the τA lobe (Figure S7D). Therefore, it is suggested that direct interaction between Pol III and TFIIIC may be not necessary for Pol III recruitment to promoter. Previous study has shown that once TFIIIB is assembled on DNA, TFIIIC is dispensable for in vitro transcription ^9^. Additionally, TFIIIC and Pol III occupancy on DNA are inversely correlated, as shown by chromatin immunoprecipitation ^27^. The interaction between Brf1, B” and τ131 also increases during transcription repression ^27^. Furthermore, smFRET data demonstrates that the DNA within the TFIIIA-TFIIIC-Brf1-TBP complex is not static and undergoes spontaneous transitions on slow time scales (10s of seconds) to a partially unwrapped state (Figure 5C). This state could be crucial for allowing Pol III to bind productively.

Finally, the TFIIIC complex needs to be displaced from DNA to allow Pol III transcription. The B” binding and completion of TFIIIB may trigger a transition to a new state where the contact between τ131 and Brf1-TBP is broken, resulting in the detachment of TFIIIC τA lobe from DNA (Figure 6, state 4). At the same time, the stable TFIIIB-DNA complex is assembled and can recruit Pol III ^9,50^. This “Pol III loading’ state may resemble yeast and human Pol III PIC structures ^15-17^. Once the PIC is formed, Pol III may initiate transcription. For transcription to occur over the full length of the gene, both TFIIIC and TFIIIA need to be displaced from the DNA.

## ACKNOWLEDGEMENTS

We thank past and present lab members for advice, assistance, and comments on the manuscript. We thank Jason Pattie for computer support. We thank Janette Meyers, Rose Marie Haynes, and Harry Scott at the PNCC for data collection support. We thank the staff at the Structural Biology Facility (SBF) of Northwestern University for technical support. Y. H. was supported by an Institutional Research Grant from the American Cancer Society (IRG-15-173-21), NIGMS (R01GM135651 and R01GM144559), NCI (P01CA092584), and an H Foundation Core Facility Pilot Project Award. R. V. was supported by NIGMS (R01GM140272) and The Searle Leadership Fund for the Life Sciences at Northwestern University. Y. H and R. V. were supported by Chicago Biomedical Consortium with support from the Searle Funds at The Chicago Community Trust. A. T. and A. R. were supported by NIGMS (5T32GM140995). A portion of this research was supported by NIH grant U24GM129547 and performed at the PNCC at OHSU and accessed through EMSL (grid.436923.9), a DOE Office of Science User Facility sponsored by the Office of Biological and Environmental Research. This work used resources of the Northwestern University Structural Biology Facility, which is generously supported by the NCI CCSG P30 CA060553 grant awarded to the Robert H. Lurie Comprehensive Cancer Center. We acknowledge the use of the Ametek K3 direct electron detector, which was generously provided by Professor Robert A. Lamb, Ph.D., Sc.D., HHMI investigator.

## AUTHOR CONTRIBUTIONS

Y He, Y Han and A Talyzina conceived the project. Y Han, S Fishbain, A Reyes, and A Talyzina purified proteins. A Talyzina and Y Han assembled the complexes and prepared cryo-EM samples. A Talyzina and Y Han processed cryo-EM data. A Talyzina built atomic models. C Banerjee and R Vafabakhsh collected and processed single molecule data. A Talyzina and Y He wrote the manuscript with input from all authors.

## DATA AVAILABILITY

Electron density map and coordinates for the TFIIIA-TFIIIC-Brf1-TBP complex bound to 5S rRNA gene have been deposited in the Electron Microscopy Data Bank (EMDB) with ID code EMDB-29071 and the Protein Data Bank (PDB) with ID code 8FFZ, respectively. Electron density map for the focused refinement on the Brf1-TBP-DNA has been deposited in the EMDB with ID code EMDB-29356. Electron density map for the TFIIIA-TFIIIC complex bound to 5S rRNA gene has been deposited in the EMDB with ID code 29358.

## FIGURE LEGENDS

**Movie S1**.

Structure of the *S. cerevisiae* TFIIIA-TFIIIC-Brf1-TBP complex bound to 5S rRNA gene.

**Movie S2**.

