## Supplementary Material for "Structural basis of TFIIIC-dependent RNA Polymerase III transcription initiation"

### TITLE

### AFFILIATIONS


### SUPPLEMENTARY FIGURE LEGENDS

**Figure S1 TFIIIA-TFIIIC complex data processing, related to Figure 1.**

(A) Cryogenic electron micrograph of TFIIIA-TFIIIC complex. (B) TFIIIA-TFIIIC cryo-electron microscopy processing pipeline. Several rounds of 3D classification resulted in a reconstruction that could be refined to an overall resolution of 6.78 Å. CTF refinement and Bayesian polishing resulted in a map with 6.62 Å resolution. (C) Fourier shell correlation (FSC) between the half maps of the final reconstruction. The reported resolution is estimated at the FSC 0.143 cut-off (dashed line). (D) Angular distribution for particles in the final reconstruction of the TFIIIA-TFIIIC complex. Color from blue to red and the height of the bars correlate with the number of particles at a specific orientation. (E) TFIIIA-TFIIIC reconstruction colored according to local resolution. The color bar from blue to red indicates the local resolution range in Å.

**Figure S2 TFIIIA-TFIIIC-Brf1-TBP complex data processing, related to Figure 1.**

(A) SDS PAGE gel of purified TFIIIA-TFIIIC-Brf1-TBP complex on DNA. Individual subunits are labeled on the gel. A band labeled with an asterisk corresponds to τ91 degradation product (B) Cryogenic electron micrograph of TFIIIA-TFIIIC-Brf1-TBP complex. (C) TFIIIA-TFIIIC-Brf1-TBP cryo-electron microscopy processing pipeline. Several rounds of 3D classification resulted in a reconstruction that could be refined to an overall resolution of 4.36 Å. Three rounds of CTF refinement and Bayesian polishing resulted in a map with 3.83 Å resolution. The Brf1-TBP-DNA part of the density was masked and separately processed in cryoSPARC. (D) Fourier shell correlation (FSC) between the half maps of the final reconstruction. The reported resolution is estimated at the FSC 0.143 cut-off (dashed line). (E) Angular distribution for particles in the final reconstruction of the TFIIIA-TFIIIC-Brf1-TBP complex. Color from blue to red and the height of the bars correlate with the number of particles at a specific orientation. (F) TFIIIA-TFIIIC-Brf1-TBP reconstruction colored according to local resolution. The color bar from blue to red indicates the local resolution range in Å. (G) Examples of model-to-map fit with several labeled bulky side chains. Model colors correspond to the colors used in the main figures.

**Figure S3 Model building details, related to Figure 1.**

(A) Alphafold structure prediction of individual domains in TFIIIA-TFIIIC-Brf1-TBP complex guided by the density map. The color bar from red (low) to blue (high) indicates the pLDDT confidence measure. (B) Brf1-TBP model on DNA was built using the Brf1-TBP-DNA model from PDB:6CNB (yellow). This model was fit as rigid body, and model fragments out of density were trimmed. (C) N-terminal TPR of τ131 in TFIIIA-TFIIIC complex was built using PDB:6YJ6. Subunit τ131 from the published model was aligned with C-terminal TPR from TFIIIA-TFIIIC-Brf1-TBP. The aligned model of τ131 from PDB:6YJ6 was fit as rigid body into the TFIIIA-TFIIIC map. Identical colors are used for τ131, τ95 and τ55 subunits in both models. (D) Comparison of possible τ131 conformations. C-terminal TPR is shown as transparent tubes. N-terminal TPR is shown as opaque tubes. The color bar from red (low) to blue (high) indicates the pLDDT confidence measure for the Alphafold prediction. The overlay of aligned τ131 models suggests how the N-terminal TPR may move relative to the C-terminal TPR. The numbers from 1 to 3 indicate the trajectory of N-terminal TPR array movement from the most closed (1) to the most open (3) conformation.

**Figure S4 TFIIIC interactions with DNA, related to Figure 3.**

(A-C) TFIIIA-TFIIIC-Brf1-TBP shown as surface and colored according to coulombic potential. The model is colored in a range from red for negative potential to blue for positive potential. Upstream (A), center (B), and downstream (C) regions of DNA are bound to positively charged patches of the protein complex. (D) Comparison of DNA-binding WH from this work (top row) with several published DNA-binding WH (bottom row). All WH domains are shown as surface and colored according to coulombic potential. The models are colored in a range from red for negative potential to blue for positive potential. (E-F) Regions of τ138 (E) and τ131 (D), which are in proximity of DNA, are colored according to the conservation score (prepared using ConSurf web server). The color bar from cyan (low) to maroon (high) indicates ConSurf sequence conservation score. DNA-contacting regions have the highest conservation score. DNA colors correspond to the colors used in the main figures.

**Figure S5 Comparison of TFIIIA ZFs, related to Figure 4.**

(A) Comparison of yeast TFIIIA with published models of *X. laevis* TFIIIA. In all cases, TFIIIA is bound to DNA, but in PDB:2HGH, TFIIIA binds RNA. ZF fold and linkers between ZF are similar between species. ZF 1 to 3 bind DNA similarly. However, the following ZFs interact with DNA in a different mode. (B) Aligned TFIIIA ZFs are colored in rainbow from N-terminus to C-terminus. Nine ZFs of *S. cerevisiae* have identical C2H2 fold, and ZF9 has a longer helix. (C) Comparison of DNA-binding TFIIIA ZF. ZF 1,2,3,5,9 bind the major grove and ZF4 binds the minor grove. (D) TFIIIA ZFs are aligned and colored according to conservation score (prepared using ConSurf web server). The color bar from cyan (low) to maroon (high) indicates ConSurf sequence conservation score. DNA-contacting ZFs have the highest conservation score on their DNA-contacting surface. DNA colors correspond to the colors used in the main figures.

**Figure S6 smFRET traces of DNA+TFIIIA+TFIIIC+Brf1-TBP, related to Figure 5.**
(A-B) single-molecule time traces of full complex, DNA+TFIIIA+TFIIIC+Brf1-TBP showing donor as a green line and acceptor as a red line. The corresponding FRET profile of the complex shows two states depicted as a blue line. (C) Similar time traces of donor (green) and acceptor (red), corresponding FRET value (blue) showing the presence of one state.

**Figure S7, TFIIIC interactions with DNA, TFIIIA, Brf1-TBP, and Pol III, related to Figure 6.**
(A) Selected 2D class averages from negative stained electron micrographs. TFIIIC lobe τB is indicated with white arrow. Each experimental condition shows either TFIIIC alone, bound to asparagine tRNA gene (DNA*), or 5S rRNA gene (DNA). (B) Side view of TFIIIA-TFIIIC-Brf1-TBP complex. (B-C) Superposition of the Pol III initial transcribing complex (PDB: 6CNB) on TFIIIA-TFIIIC-Brf1-TBP showed severe clashes between Pol III (grey surface) and TFIIIC (C). However, detachment of Brf1-TBP-DNA from TFIIIC surface may result in simultaneous binding of Pol III PIC and TFIIIA-TFIIIC complex to DNA (D). The direction of potential Pol III PIC and DNA displacement is indicated with an arrow.

### METHODS

#### Purification of TFIIIA

Two liters of transformed BL21 (DE3) pRARE cells were grown in LB at 37 °C to an OD600 of 0.6. Then 50 µM $ZnSO_4$ was added, the cells were induced with 1 mM IPTG, and the protein was expressed for two hours at 37 °C. Cells were pelleted for 20 minutes at 4000 x g and resuspended in 40 mL of buffer A (20mM HEPES 7.6, 250 mM NaCl, 5 mM $MgCl_2$, 50 µM ZnSO4, 10% glycerol, 5mM dithiothreitol [DTT], 1 mM phenylmethylsulfonyl fluoride [PMSF]). After cell lysis by sonication, cell pellet was collected by centrifugation for 10 minutes at 4000 x g. The pellet was resuspended in buffer A + 5M urea, then briefly sonicated again and incubated on nutator overnight at 4 °C. Next day, cell debris was removed by centrifugation at 15000 x g at 4 °C for 30 minutes and the supernatant was filtered. The supernatant was loaded twice onto a gravity column with 1 ml of HIS-Select Nickel Affinity Gel (Sigma-Aldrich) equilibrated with buffer A + 5M urea + 20mM imidazole. The resin was washed six times with 2 ml of buffer A + 5M urea + 20mM imidazole. The protein was eluted in 1 ml fractions using buffer A + 5M urea + 300mM imidazole. Fractions containing TFIIIA were pooled together and diluted in three steps over 45 minutes with Buffer A + 1mM PMSF + 5mM DTT. The diluted protein was dialyzed against 1L of Buffer A + 1mM PMSF + 10mM BME in 3.5kDa cut-off snakeskin tubing (Thermo Fisher) overnight at 4 °C. Next day aggregation was removed by centrifugation at 10000 x g for 5 minutes. Supernatant containing refolded TFIIIA was flash-frozen in liquid nitrogen.

#### Purification of TFIIIC

TFIIIC was purified from an *S. cerevisiae* strain with a TAP tag at the C terminus of τ60 (GE Dharmacon, YSC1178-202233621). Eight liters of yeast were grown overnight. Cells were harvested by centrifugation and resuspended in 200ml of cold TAP extraction buffer (40 mM HEPES pH 8, 250 mM ammonium sulfate, 1 mM EDTA, 10% glycerol, 0.1% Tween-20, 1 mM DTT, 1 mM PMSF, 2mM benzamidine, 0.3 µg/ml leupeptin, 1.4 µg/ml pepstatin, 2 µg/ml chymostatin). Cells were lysed using BeadBeater (Biospec Products). Cell debris was removed by centrifugation at 15000 x g at 4 °C for two hours. The lysate was incubated with 2ml of IgG Sepharose beads (GE Healthcare) for two hours at 4 °C. The beads were washed and resuspended in 4ml of cold TEV cleavage buffer (10 mM HEPES pH 8, 200 mM NaCl, 0.1% NP-40, 0.5 mM EDTA, 10% glycerol). TEV cleavage was performed using 25 µg of TEV protease at room temperature (RT) for one hour. TEV flowthrough was collected and $CaCl_2$ was added for a final concentration of 2 mM. 800 µl of Calmodulin Affinity Resin (Agilent Technologies) was washed with Calmodulin binding buffer (15 mM HEPES pH 8, 1 mM magnesium acetate, 1 mM imidazole, 2 mM CaCl2, 0.1% NP-40, 10% glycerol, 200 mM ammonium sulfate, 1 mM DTT, 1 mM PMSF, 2mM benzamidine, 0.3 µg/ml leupeptin, 1.4 µg/ml pepstatin, 2 µg/ml chymostatin) and incubated with TEV flowthrough overnight at 4 °C. Following incubation, the beads were washed with Calmodulin binding buffer, followed by Calmodulin wash buffer (same as the binding buffer but with 0.05% NP-40), followed by Calmodulin transfer buffer (same as wash buffer but without $CaCl_2$). 400 µl of Calmodulin elution buffer (15 mM HEPES pH 8, 1 mM magnesium acetate, 1 mM imidazole, 5 mM EGTA, 10% glycerol, 0.05% NP-40, 200 mM ammonium sulfate) was added to the beads and incubated for 45 minutes at 4 °C. First, 400 µl fraction was eluted, another 400 µl of elution buffer was added to the beads, and eluted after 5 minutes. The following fractions were eluted immediately. Fractions containing protein were pooled together, concentrated and flash-frozen in liquid nitrogen.

**Purification of Brf1-TBP**

The chimera protein Brf1N-TBPc-Brf1C [1] was purified as follows. Two liters of transformed BL21 (DE3) pRARE cells were grown in LB at 37 °C to an OD600 of 0.6, the cells were induced using 0.5 mM IPTG, and the protein was expressed overnight at 18 °C. Cells were harvested by centrifugation and resuspended in 35 ml of BT lysis buffer (20 mM HEPES pH 7.6, 25 µM EDTA, 1.14 M NaCl, 5% glycerol, 10 mM BME, 0.5 mM PMSF, 1 µg/ml leupeptin, 1 µg/ml pepstatin, 300 µg/ml lysozyme), followed by incubation on ice for one hour and followed by sonication. Cell debris was removed by centrifugation at 13000 x g at 4 °C for one hour. The lysate was loaded onto a gravity column with 500 µl of HIS-Select Nickel Affinity Gel (Sigma-Aldrich) equilibrated with BT lysis buffer. The resin was washed five times with 1 ml of BT wash buffer 1 (20 mM HEPES pH 7.6, 7 mM MgCl2, 0.5 M NaCl, 10 mM imidazole, 5% glycerol, 10 mM BME, 0.5 mM PMSF, 1 µg/ml leupeptin, 1 µg/ml pepstatin) and five times with 1 ml of BT wash buffer 2 (20 mM HEPES pH 7.6, 7 mM MgCl2, 0.5 M NaCl, 20 mM imidazole, 5% glycerol, 10 mM BME, 0.5 mM PMSF, 1 µg/ml leupeptin, 1 µg/ml pepstatin). Protein was eluted in five 1 ml fractions with BT elution buffer (20 mM HEPES pH 7.6, 7 mM MgCl2, 0.5 M NaCl, 200 mM imidazole, 5% glycerol, 10 mM BME, 0.5 mM PMSF, 1 µg/ml leupeptin, 1 µg/ml pepstatin). Fractions containing Brf1-TBP were pooled and dialyzed against 500 ml of dialysis buffer (20mM HEPES pH 7.6, 200mM NaCl, 7mM MgCl2, 0.01% Tween-20, 10% glycerol, 0.2mM PMSF, 10mM BME) in 12 kDa Pur-A-Lyzer Maxi (SigmaAldrich) overnight at 4 °C.

Next day the dialysis buffer was replaced with fresh dialysis buffer and the protein was dialyzed for another 5 hours. Fractions containing protein were pooled together and flash-frozen in liquid nitrogen.

**Complex assembly**

First, 2 pmol of 5S rRNA gene DNA template (sense: 5'-/5BiotinTEG/TACGGACCATGGAATTCCCCAGT AACATGTCTGGACCCTGCCCTCATATCACCTGCGTTTCCGTTAAACTATCGGTTGCGG CCATATCTACCAGAAAGCACCGTTTCCCGTCCGATCAACTGTAGTTAAGCTGGTAAG AGCCTGACCGAGTAGTGTAGTGGGTGACCATACGCGAAACTCAGGTGCTGCAATCT -3', antisense: 5'-AGATTGCAGCACCTGAGTTTCGCGTATGGTCACCCACTACACTACTCGGTCAGGCTC TTACCAGCTTAACTACAGTTGATCGGACGGGAAACGGTGCTTTCTGGTAGATATGGC CGCAACCGATAGTTTAACGGAAACGCAGGTGATATGAGGGCAGGGTCCAGACATGT TACTGGGGAATTCCATGGTCCGTA -3') was mixed with 100 nmol TFIIIA and incubated at RT for 5 minutes. Then 200 nmol TFIIIC was added and incubated at RT for 5 minutes. For the TFIIIA-TFIIIC-Brf1-TBP complex, the previous step was followed by the addition of 150 nmol Brf1-TBP. The salt concentration was adjusted to 100 mM KCl with the addition of buffer 1 (12 mM HEPES pH 7.6, 0.12 mM EDTA, 12% glycerol, 8.25 mM $MgCl_2$, 1 mM DTT, and 0.05% NP-40). All components were incubated for an additional 5 minutes at RT before binding to T1 streptavidin beads (Fisher Scientific) at RT for 15 minutes. Assembled complexes were washed with buffer 2 (10 mM HEPES pH 7.6, 10 mM Tris pH 7.6, 5% glycerol, 5 mM $MgCl_2$, 50 mM KCl, 1 mM DTT, and 0.05% NP-40) and eluted with buffer 3 (10 mM HEPES pH 7.6, 5% glycerol, 10 mM $MgCl_2$, 50 mM KCl, 1 mM DTT, 0.05% NP-40, and 30 units EcoRI-HF (New England Biolabs).

**Negative stain EM data collection and processing**

Negative stain samples were prepared using 400 mesh copper grids (Electron Microscopy Sciences) coated with continuous carbon on a nitrocellulose support film. Before usage, they were glow-discharged for 10 seconds with 25 W of power using the Solarus plasma cleaner 950 (Gatan). Purified TFIIIA-TFIIIC and TFIIIA-TFIIIC-Brf1-TBP complexes in buffer 4 were cross-linked with 0.05% glutaraldehyde for 10 minutes on ice and incubated for 10 minutes on a grid in a homemade humidity chamber at 4 °C. The grid was stained on four 40 µL drops of 2% uranyl formate solution for 5, 10, 15, and 20 seconds sequentially and blotted dry with #1 filter paper (Whatman). Images were collected on a Jeol 1400 microscope equipped with a Gatan 4k × 4x CCD camera at 30,000x magnification (3.71 Å/pixel), a defocus range of -1.5 to -3 µm, and 20 $e^-/Å^2$ total electron dose using Leginon [2].

Particles were picked using DogPicker, extracted, and 2D classified using iterative MSA/MRA topological alignment within the Appion data processing software [3-6]. A particle stack of ~50,000 particles with a box size of 96 x 96 pixels was subjected to iterative, multi-reference projection-matching 3D refinement using EMAN2 software package to generate an initial reference for cryo-EM data processing [7].

### Cryo-EM sample preparation

Cryo-EM samples were prepared using Quantifoil 2/1 300 mesh copper grids (EMS). Grids were glow discharged for 10 seconds with 25 W of power using the Solarus plasma cleaner 950 (Gatan), and then a thin layer of graphene oxide was applied as described previously [8]. Purified TFIIIA-TFIIIC and TFIIIA-TFIIIC-Brf1-TBP samples (~3.5 μL) were incubated with 0.05% glutaraldehyde for 10 minutes on ice. The sample was applied to a grid in a Vitrobot Mark IV (Thermo Fisher Scientific) operating at 4 °C with 100% humidity. After 5 minutes of incubation, the sample was blotted with 10 force for four seconds and immediately plunged into liquid ethane cooled to liquid nitrogen temperature.

### Cryo-EM data collection and processing

Cryo-EM data were collected at the Pacific Northwestern Center for Cryo-EM (PNCC). Images were collected using semi-automated data collection in Serial EM [9] on a Titan Krios transmission electron microscope (TEM) operated at 300 keV (Thermo Fisher Scientific), equipped with a Quantum energy filter (Gatan), and with a K3 direct detector (Gatan) operating in super-resolution mode.

For the TFIIIA-TFIIIC sample, images were collected at a magnification of 30,000X (0.5395 Å/pixel) using a defocus range of -1.5 to -3 μm with a dose rate of 1 $e^-$/pixel/frame for a total dose of 50 $e^-/Å^2$. A dataset of 11,645 images was collected. For the TFIIIA-TFIIIC-Brf1-TBP sample, images were collected at a magnification of 30,000X (0.528 Å/pixel) using a defocus range of -2 to -5 μm with a dose rate of 1 $e^-$/pixel/frame for a total dose of 60 $e^-/Å^2$. A dataset of 23,211 images was collected.

RELION 3.1 was used for all pre-processing, 3D classification, model refinement, post-processing, and local-resolution estimation jobs [10]. Particles were picked using Gautomatch (developed by K. Zhang, MRC Laboratory of Molecular Biology, Cambridge, UK) and Laplacian-of-Gaussian (LoG) picking in RELION 3.1. Duplicated particles were removed. Local CTF of each micrograph was determined using CTFFIND-4.1 [5].

For the TFIIIA-TFIIIC complex, a stack of 2,176,308 particles was binned by a factor of 2 (2.158 Å/pixel) and extracted with a box size of 132 pixels. First round of 3D classification (10 classes) with the negative stain reconstruction as a reference was used to clean up the particle stack. One class of 159,078 particles showed clear structural features of the TFIIIA-TFIIIC complex and was selected for further processing. The selected particles were 3D auto-refined and used in the second round of 3D classification (5 classes). Three classes (109,548) were selected and 3D auto-refined. Another round of 3D auto-refinement was performed with a soft mask, resulting in a 6.78 Å resolution reconstruction. All reported resolutions correspond to the gold-standard Fourier shell correlation (FSC) using the 0.143 criteria [11]. Three rounds of per-particle CTF refinement were performed, followed by Bayesian particle polishing. 3D auto-refinement using the polished particles resulted in a 6.62 Å resolution map.

For the TFIIIA-TFIIIC-Brf1-TBP complex, a stack of 5,748,589 particles was binned by a factor of 2 (2.112 Å/pixel) and extracted with a box size of 132 pixels. First round of 3D classification (10 classes) with the negative stain reconstruction as a reference was used to clean up the particle stack. One class of 665,210 particles showed clear structural features of the TFIIIA-TFIIIC-Brf1-TBP complex and was selected for further processing. The selected particles were 3D auto-refined and used in the second round of 3D classification (3 classes) without alignment. One class (78,512 particles) was selected. The selected particles were 3D auto-refined, re-centered, and re-extracted without binning (1.056 Å/pixel, box size = 264 pixels). Another round of 3D auto-refinement was performed with a soft mask applied around the whole complex, resulting in a 4.36 Å resolution reconstruction. The particle stack was re-extracted with a bigger box size (384 pixels). Three rounds of 3D auto-refinement, per-particle CTF refinement and Bayesian particle polishing were performed. 3D auto-refinement using the polished particles resulted in a 3.83 Å resolution map. To improve the map quality of the Brf1-TBP part, focused 3D variability analysis was performed in cryoSPARC [12]. Briefly, the particle stack that resulted in 4.36 Å resolution reconstruction, was imported into cryoSPARC, where it was used for Non-uniform (NU) refinement, local refinement (soft mask applied to focus on Brf1-TBP-DNA density), signal subtraction and 3D variability analysis in cluster mode. This procedure produced a 7.14 Å resolution reconstruction of Brf1-TBP-DNA density. In parallel with post-processing done in RELION 3.1, DeepEMhancer was used to better correct local B-factors and produced cleaner maps for model building and docking [13].

**Model building and refinement**

TFIIIA-TFIIIC-Brf1-TBP model:

The resolution of the TFIIIA-TFIIIC-Brf1-TBP complex map (3.83 Å) allowed for AlphaFold-guided model building for the entire complex [14]. Alphafold was used to predict individual domains or full subunits of TFIIIA and TFIIIC, and the produced models were fitted as a rigid body into the density map using UCSF Chimera [15]. Published experimental structures were used in rigid body fitting as well. Manual adjustments were made in Coot [16] and ISOLDE [17]. The final model was refined in Phenix. The refined model was inspected with the help of ISOLDE, where clashes and rotamer outliers were resolved.

TFIIIA:

*X. laevis* structure of ZF 1 to 3 bound to DNA (PDB:1TF3) was fitted as rigid body into the TFIIIA density. *S. cerevisiae* TFIIIA Alphafold prediction was aligned with the experimental *X. laevis* structure, and individual ZF was fitted as rigid body into the TFIIIA density. Predicted regions that were not visible in the density were trimmed.

TFIIIC:

The structure of the subcomplex of τ60 and τ91 (PDB: 2J04) was fitted as rigid body into the density. *S. cerevisiae* Alphafold predictions for τ60 and τ91 were aligned with the experimental structure, and flexible loops, not visible in the density, were trimmed. The cryo-EM structure of the τA lobe comprised of τ131, τ95 and τ55 subunits (PDB: 6YJ6) was fitted as rigid body. Alphafold predictions were aligned with subunits and manually adjusted. Predicted regions that were not visible in the density were trimmed. For τ138, individual folded domains from Alphafold prediction were fitted as rigid body into the map, and connecting linkers were manually adjusted.

Brf1-TBP:

Brf1-TBP model from *S. cerevisiae* Pol III PIC structure (PDB:6CNB) was fitted into Brf1-TBP density as rigid body, and flexibly tethered domains of Brf1 that could not be traced in the density (cyclin fold 1 and zinc ribbon), were trimmed. This model was not manually adjusted due to low resolution of the map in this area.

DNA:

Double strand DNA model with 5S rRNA gene sequence was generated using webserver (http://www.scfbio-iitd.res.in/software/drugdesign/bdna.jsp). The DNA register was traced using sequence alignment between *X. laevis* and *S. cerevisiae* ICR and structure of *X. laevis* structure of ZF 1 to 3 bound to DNA (PDB:1TF3), as well as the positioning of Brf1-TBP at the upstream region. The DNA was fitted into the density in ISOLDE.

TFIIIA-TFIIIC model:

The model of the TFIIIA-TFIIIC-Brf1-TBP complex was fitted in the density map of the TFIIIA-TFIIIC complex. TFIIIC subunits τ55, τ95, τ91, and τ60 completely fit in the map. Protein-DNA contacts within the τB model are identical as well. C-terminal TPR and helical domains of τ131 are placed identically to the TFIIIA-TFIIIC-Brf1-TBP complex. N-terminal TPR cannot be resolved, but low-resolution features suggested a conformation of τ131, similar to the one in τA lobe structure (PDB: 6YJ6). TFIIIA zinc fingers 1-5 interact with DNA, while zinc fingers 6-8 protrude away from the DNA. The overall position of TFIIIA is identical to the TFIIIA-TFIIIC-Brf1-TBP complex; however, zinc finger 9 and adjacent helix are not visible in the map. DNA in this complex is also positioned identically to the TFIIIA-TFIIIC-Brf1-TBP complex.

UCSF Chimera and UCSF Chimera X were used for figure and movie generation [15,18]. ConSurf web server was used for estimating the evolutionary conservation of residues [19].

**Preparation of DNA constructs for smFRET**

The PAGE-purified DNA oligonucleotides containing biotin for immobilization or amine modifications were purchased from Genscript (sense: AACATGTCTGGACCCTGCCCTCATATCACCTGCGTTTCCGTTAAACTATCGGTTGCGG CCATATCTACCAGAAAGCACCGTTTCCCGTCCGATCAACTGTAGTTAAGCTGGTAAG AGCCTGACCGAGTAGTGTAGTGGG/iAmMC6dT/GACCATACGCGAAACTCAGGTGCT GCAATCTGTAGATTCATTGGACTGGTG; antisense: AGATTGCAGCACCTGAGTTTCGCGTATGGTCACCCACTACACTACTCGGTCAGGCTC TTACCAGCTTAACTACAGTTGATCGGACGGGAAACGGTGCTTTCTGGTAGATA/iAmM C6dT/GGCCGCAACCGATAGTTTAACGGAAACGCAGGTGATATGAGGGCAGGGTCCA GACATGTTACTGGGGAATTCCATGGTCCGTA;
sense, biotinylated: 5’-/5BiotinTEG/TACGGACCATGGAATTCCCCAGT-3’).
Oligonucleotides, carrying an amine group, were labeled with NHS ester-conjugated fluorescent dyes. For the labeling reaction, 6.25 μl of 40 μM amine-modified antisense DNA was mixed

with an excess of NHS ester-conjugated Cy3 dye (200:1 ratio of dye to DNA) in 100 mM Na2B4O7 pH8.4 and 40% DMSO. The mix was incubated at 25°C for 6 hours, then at 4°C overnight on a gently shaking mixer in the dark. The excess dye was removed by ethanol precipitation and 70% ethanol wash. The labeling reaction was performed twice to increase labeling efficiency. Amine-modified sense DNA was labeled with NHS ester-conjugated Cy5 dye in the same way. The dsDNA construct (1 μM) was prepared by mixing the biotinylated sense DNA with sense DNA-Cy5 and antisense DNA-Cy3 at a molar ratio of 1.2:1.1:1 and annealed in water in a heat block by incubation at 95°C for 5 minutes followed by gradual cooling to RT.

**Complex assembly and pulldown for smFRET**

To achieve consistency between the two methods, we performed the protein-DNA complex assembly in the same way as preparing the cryo-EM samples with a few modifications. The annealed and Cy3-/Cy5-labeled dsDNA construct was used instead of the 5S rRNA gene DNA template. After the complex elution with EcoRI, 1.2 pmol of antisense biotinylated DNA (5'-/5BiotinTEG/CACCAGTCCAATGAATCTAC-3') was added to the elution and incubated at RT for 10 minutes. The integrity of TFIIIA-TFIIIC and TFIIIA-TFIIIC-Brf1-TBP complexes assembled on fluorescently labeled DNA was confirmed by negative stain EM. The sample was incubated on ice before the smFRET experiment.

**smFRET measurements**

Single-molecule experiments were performed in a flow chamber prepared by sandwiching PEG (mPEG and biotin-PEG 1% (w/w), Laysan Bio) passivated glass coverslip (VWR) and slides (ThermoFisher Scientific) [20]. Before imaging, flow chambers were washed with T-50 buffer (50 mM NaCl, 10 mM Tris, pH 7.4) and incubated with 50 nM NeutrAvidin (ThermoFisher Scientific) for 2 minutes. Then the flow chambers were washed with T-50 to remove unbound NeutrAvidin. Finally, the pre-assembled complex of labeled DNA and proteins, as described above, was injected into the chamber to immobilize on the surface. The complex was diluted and imaged in an imaging buffer consisting of 4 mM Trolox, 10 mM HEPES, 10 mM $MgCl_2$, 50 mM KCl, 5uM $ZnCl_2$, 5 % glycerol, and an oxygen-scavenging system consisting of 4 mM protocatechuic acid (Sigma) and 1.6 $Uml^{-1}$ bacterial protocatechuate 3,4-dioxygenase (rPCO) (Oriental Yeast), pH 7.35. All the chemicals were purchased from Sigma (purity >99.9%). smFRET data were recorded at 50 ms time resolution using a TIRF microscope with a 100x oil-immersion objective (Olympus, NA 1.49). The donor and acceptor fluorophores were excited using 532 nm and 638 nm lasers, respectively.

**smFRET data analysis**

Single-molecule fluorescence intensity traces were analyzed using smCamera (http://ha.med.jhmi.edu/resources/), custom-written Matlab (MathWorks) scripts, and OriginPro. smFRET particles were selected based on the Gaussian intensity profile of the spots, acceptor brightness at least 5% above the background, and the acceptor signal upon donor excitation. The donor and acceptor intensity traces were selected based on the following criteria. A single donor and a single acceptor bleaching step during the acquisition time window, stable total intensity ($I_D$

+ $I_A$), and anticorrelated intensity profiles of donor and acceptor without blinking. Two authors independently examined all the data and found identical results. Each experiment, unless otherwise noted, was carried out three times to ensure that the findings could be repeated. Apparent FRET efficiency was calculated using $(I_A - 0.088 \times I_D)/(I_D + I_A)$, where $I_D$ and $I_A$ are raw donor and acceptor intensities, respectively. smFRET histograms for the DNA, DNA+TFIIIA+TFIIIC, and DNA+TFIIIA+TFIIIC+Brf1-TBP were generated from the 3, 4, and 16 movies from different days.
To determine the FRET value, the smFRET histograms were fitted using OriginPro with 1 or 2 Gaussian distributions.

$$y(x) = \sum_{i=1}^{n} \frac{A_i}{w_i\sqrt{\frac{\pi}{2}}} e^{-2\frac{(x-xc_i)^2}{w_i^2}}$$

where $n$ is the number of Gaussians, $A$ is the peak area, $xc$ is the FRET peak center, and $w$ is the full-width half maximum for each peak.
To derive the transition density plots (TDPs), all the time traces were idealized by fitting with hidden Markov model (HMM) using vbFRET software in Matlab [21]. Traces were fit with 1 to 4 step models and majority of traces fit with a 1 or 2 steps. Then, from the idealized traces, all the transitions were extracted to create a transition density plot.
